# Distinct population dimensions separate sensory adaptation and expectation-related signals in visual cortex

**DOI:** 10.64898/2026.09.30.755465

**Authors:** Soyoun Kim, Arenski Vazquez, Renata Batista-Brito, Lucas Sjulson

## Abstract

Mismatch negativity is conventionally quantified as a scalar response-amplitude difference between rare deviant and repeated standard stimuli, leaving its organization at the neuronal population level unresolved. We recorded population activity in mouse primary visual cortex (V1) during visual odd-ball and control paradigms while testing contributions from anterior cingulate cortex (ACC) input using optogenetic inhibition. Mismatch modulated firing rates bidirectionally, and these changes largely canceled out on average, yet produced robustly separable population activity patterns. On the stimulus-encoding manifold, repeated standard exposure produced standard-specific response attenuation and a shared rotation of standard and deviant representations, revealing a component not isolated by conventional stimulus-specific adaptation and deviance-detection contrasts. Off the manifold, standards and deviants occupied opposing positions along an ACC-modulated axis consistent with expectation confirmation and violation. Thus, visual mismatch responses engage separable transformations of stimulus representations and expectation-related population activity, with ACC input selectively contributing to the latter while not significantly affecting stimulus encoding or on-manifold adaptation.

## Introduction

Mismatch negativity (MMN) is an electrophysiological response evoked when a rare “deviant” sensory stimulus occurs among frequently repeated “standard” stimuli. First characterized in the auditory system, MMN has been widely studied as a marker of sensitivity to sensory regularities and their violation (1–3). Related mismatch (MM) responses have also been observed in the visual system, suggesting that recent stimulus history influences cortical processing across modalities (4–8). Because MMN is typically measured using electroencephalography (EEG) or local field potentials (LFPs), it provides an aggregate measure of activity across heterogeneous neuronal populations. However, these aggregate signals do not directly reveal how individual neurons or neuronal populations contribute to mismatch responses. Here, we use MM responses more broadly to describe differences in unit and population activity associated with the presentation of standards and deviants.

MM responses combine at least two effects: repeated presentation of the standard attenuates sensory responses, whereas a rare deviant can recruit mechanisms related to the detection of a violated regularity. To distinguish these contributions, many studies use a control condition in which multiple stimuli are presented in random order, so that no stimulus serves as a repeated standard (9, 10). Thus, a physically identical stimulus can be compared across standard, deviant, and control conditions, with the deviant and control matched in presentation probability. The difference between control and standard responses (*C − S*) is commonly used as an operational measure of stimulus-specific adaptation (SSA), whereas the difference between deviant and control responses (*D − C*) is used as an operational measure of deviance detection (DD). SSA is generally interpreted as a reduction in sensory responsiveness resulting from repeated exposure to specific stimulus features. DD is interpreted as expectation-related because deviants and controls are matched in probability but differ in whether they violate the dominant sequence regularity. However, interpreting these contrasts as distinct mechanisms requires assumptions about how stimulus history influences sensory responses. In particular, repeated standard exposure may also alter responses to deviants, introducing adaptation-related contributions into the *D − C* contrast. Moreover, the *C − S* contrast may capture changes in sensory processing beyond standard-specific response attenuation. Consequently, these scalar contrasts do not necessarily isolate independent neuronal mechanisms. We therefore distinguish these operational contrasts from the mechanisms they are intended to estimate.

Decomposing MM responses using scalar response-amplitude contrasts has been valuable, particularly because much previous work has relied on EEG, LFP, or single-unit activity, for which amplitude is a natural summary statistic. However, neuronal population activity can reveal structure that is not captured by scalar response measures, including how sensory information is organized within high-dimensional neural state spaces. Applied to MM processing, a population-level approach can determine whether adaptation and expectation-related activity alter the same stimulus-encoding dimensions, recruit distinct population dimensions, or transform the geometry of stimulus representations. Importantly, joint analysis of standard, deviant, and control responses can also reveal transformations shared by standard and deviant representations that may contribute to both conventional contrasts without being independently identified by either. Moreover, population patterns preserve opposing changes across neurons that may largely cancel when activity is reduced to an average response. By resolving these distinct patterns of population activity, neural state-space analyses can reveal how MM-related transformations are organized within and beyond stimulus-encoding dimensions. This perspective is especially relevant in the visual cortex, where stimulus information is represented through structured population activity (11, 12).

To examine visual MM at the population level, we performed 64-channel silicon probe recordings in mouse V1 while head-fixed animals passively viewed drifting gratings in oddball and control paradigms. We compared responses to the same stimulus orientation when presented as a standard, deviant, or control. MM effects modulated firing rate bidirectionally across units and largely canceled in aggregate, yet produced distinguishable population patterns comprising three geometrically separable components. First, repeated standard exposure rotated both standard and deviant representations on the stimulus-encoding manifold, revealing a transformation shared across the two conditions that is not independently isolated by conventional SSA and DD contrasts. We term this component shared adaptation. Second, standard responses were attenuated on-manifold, consistent with sensory adaptation and more closely aligned with the conventional SSA contrast. Third, standards and deviants occupied opposing positions along an off-manifold, expectation-related axis, consistent with expectation confirmation and violation, respectively. ACC inhibition contracted both poles toward the control response, indicating that ACC input contributes to this bidirectional population signal. Together, these findings reveal a population-level organization of visual MM that distinguishes shared adaptation, on-manifold standard-specific attenuation, and off-manifold expectation-related activity. These results demonstrate that visual MM involves distinct transformations of sensory representations and expectation-related population activity, with ACC input selectively contributing to the latter.

## Results

### V1 population response patterns encode visual mis-match beyond average response amplitude

To examine how stimulus repetition and probability shape visual responses in V1, we presented drifting gratings of different orientations in oddball and control paradigms while recording neural activity with a 64-channel silicon probe (Figs. 1A,B). In each oddball block, one orientation occurred frequently as the standard and an orthogonal orientation occurred rarely as the deviant; their roles were reversed in a flip-flop block. In the control block, eight orientations were presented with equal probability in a randomized sequence. The same stimulus orientation could therefore be compared when presented as a standard, deviant, or control. In the conventional scalar framework, *C − S* is termed the SSA contrast, *D − C* is termed the DD contrast, and *D − S* is the MM contrast (Fig. 1C). These labels describe operational comparisons; whether they uniquely isolate distinct mechanisms is addressed below. LFP responses provided a conventional aggregate reference (Fig. S1), whereas our primary analyses examined single-unit and population activity. Because arousal and ongoing behavior influence V1 population activity (11, 13), we compared pupil size and facial motion across stimulus conditions in animals with synchronized video recordings. We detected no significant differences in either measure among standard, control, and deviant trials (Fig. S2A,B), suggesting that differences in stimulus condition were not associated with detectable changes in these measured behavioral-state variables.

**Fig 1.**
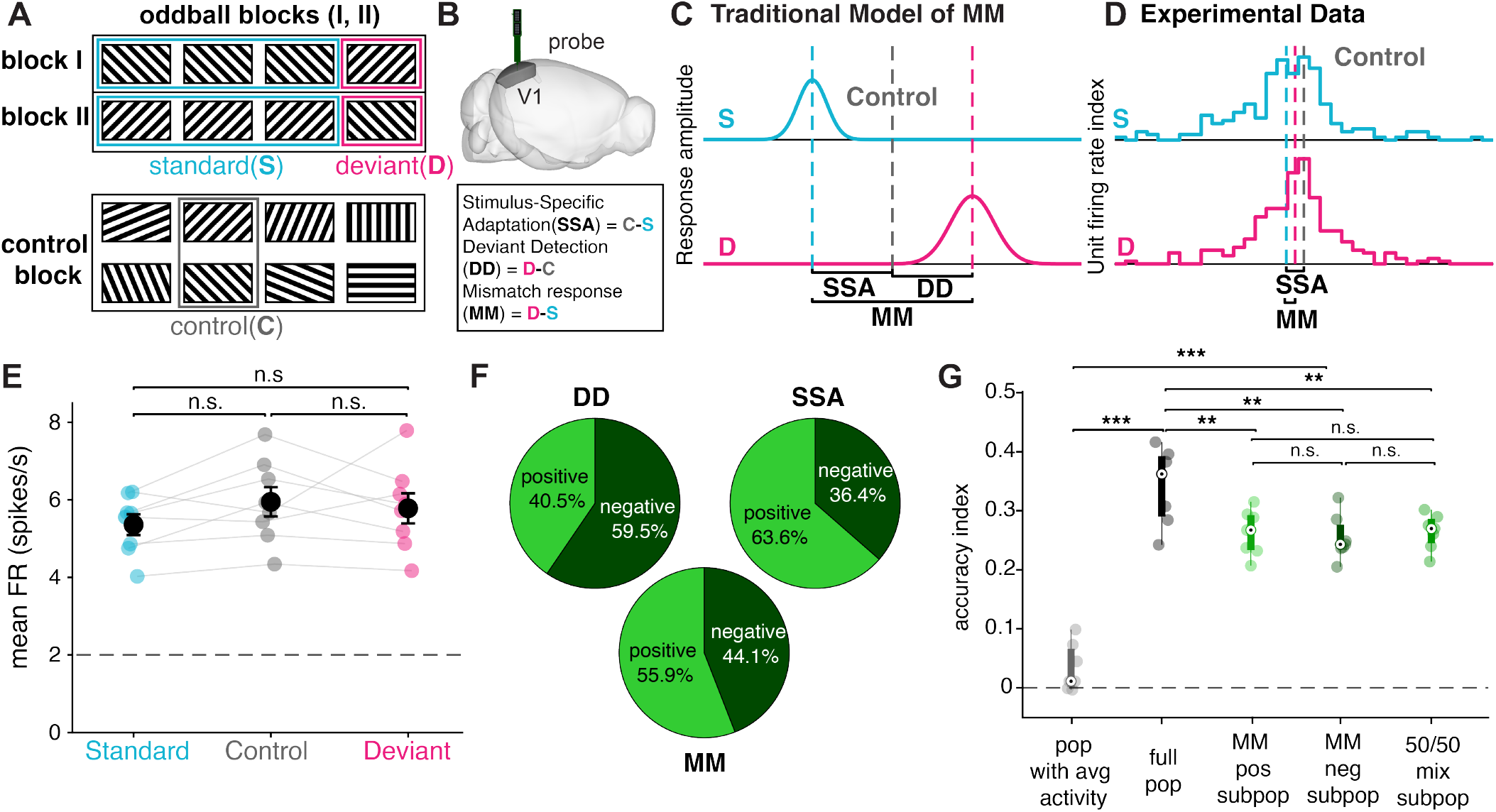
V1 population patterns encode visual mismatch beyond average response amplitude. **A**. Visual oddball and control paradigms. Responses were compared for the same stimulus orientation presented as a standard, deviant, or control. **B**. Acute silicon-probe recordings were performed in mouse V1. **C**. Conventional scalar contrasts: *C − S* is termed SSA, *D − C* is termed DD, and *D − S* is the MM contrast. **D**. Individual-unit firing rates in a representative session were modulated bidirectionally, with minimal change in average firing rate relative to held-out control responses. **E**. Across sessions, population-averaged responses did not differ significantly among conditions (*n* = 7 sessions; paired *t*-tests: D versus C, *p* = 0.657; C versus S, *p* = 0.0577; D versus S, *p* = 0.0746). **F**. Fractions of units with positive and negative *C − S, D − C*, and *D − S* modulation, classified by contrast sign. Substantial fractions were modulated in both directions. **G**. Cross-validated standard-versus-deviant decoding using population-average firing rate, the full population activity pattern, MM-positive [FR(*D*) *>* FR(*S*)] or MM-negative [FR(*D*) *<* FR(*S*)] subpopulations, or a size-matched mixed subpopulation containing equal numbers of MM-positive and MM-negative units. All pattern-based decoders exceeded shuffled performance, whereas average firing rate did not (one-sample *t*-tests: average, *p* = 0.0686; full, *p* = 8.27 *×* 10^*−*6^; positive, *p* = 1.98 *×* 10^*−*6^; negative, *p* = 2.15 *×* 10^*−*6^; mixed, *p* = 3.98 *×* 10^*−*7^). The full decoder contained the most units and thus achieved the highest accuracy (full versus positive, *p* = 0.00224; full versus negative, *p* = 0.00599; full versus mixed, *p* = 0.00213). MM-positive and MM-negative subpopulations performed comparably (*p* = 0.643), and the mixed subpopulation did not outperform either group (positive, *p* = 0.848; negative, *p* = 0.415), suggesting that positive and negative modulation carried similar rather than complementary MM information.

In a representative recording, both standard and deviant stimuli evoked roughly equal population-averaged firing rates to their matched controls (Fig. 1D), with mean responses not differing significantly across sessions (Fig. 1E). This weak aggregate effect did not reflect an absence of MM modulation: individual units showed substantial changes in both directions for the *C − S, D − C*, and *D − S* contrasts, with no predominance of changes in the direction predicted by the conventional response-amplitude model (“positive,” Fig. 1F). Thus, rather than uniformly decreasing for standards and increasing for deviants, V1 unit responses were heterogeneous and largely canceled in aggregate. Accordingly, decoding based only on population-averaged firing rate carried little information about whether a stimulus was standard or deviant (Fig. 1G). In contrast, decoders based on population activity patterns reliably distinguished the two conditions. Positively and negatively modulated units produced comparable decoding accuracy, indicating that modulation in either direction was similarly informative. Moreover, a size-matched subpopulation containing equal proportions of positively and negatively modulated units did not produce superior decoding performance, suggesting that positive and negative modulation do not encode different or complementary types of information. Visual MM was therefore encoded by patterns of activity across units and was poorly captured by a single axis of average response amplitude.

### Repeated standard exposure rotates representations on the stimulus-encoding manifold

We next examined how repeated standard exposure altered the geometry of stimulus representations. The many-standards control provided a reference because it included eight stimulus orientations, each presented with the same probability as the deviant stimulus in the oddball blocks. By applying principal component analysis to the trial-averaged population responses for all eight control orientations, we identified a seven-dimensional subspace that we define as the stimulus-encoding manifold. The first two principal components provided a visualization in which control orientations followed distinguishable trajectories, whereas all quantitative analyses used the full seven-dimensional manifold (Fig. 2A). We next projected the standard and deviant responses into this fixed, control-defined reference space (Fig. 2B), allowing us to compare their population representations with those evoked by the same stimulus orientation in the control condition.

**Fig 2.**
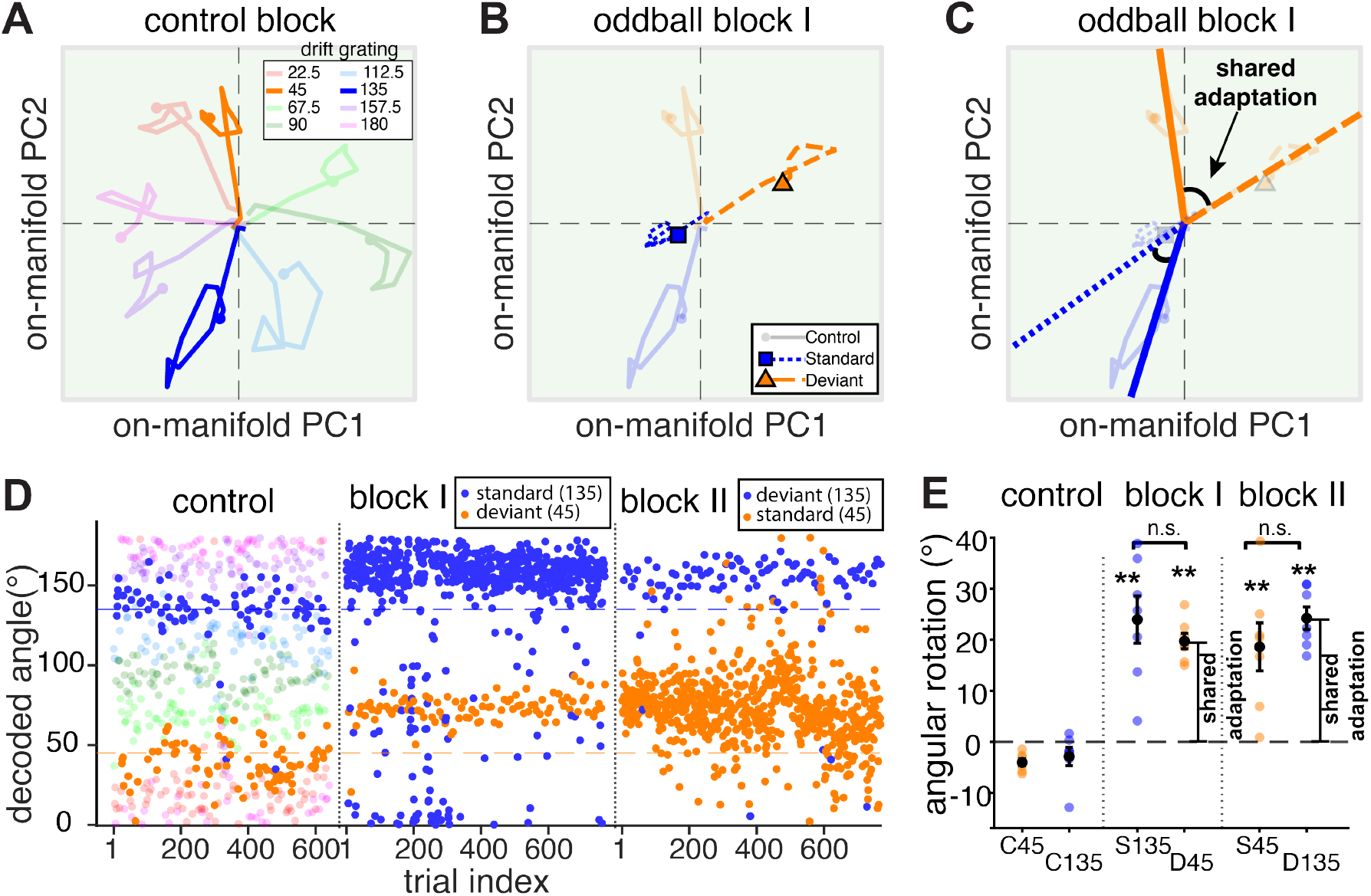
Repeated standard exposure rotates representations on the stimulus-encoding manifold. **A**. Trial-averaged responses to eight control orientations defined a seven-dimensional stimulus-encoding manifold. The first two principal components are shown for visualization; colored trajectories depict time-resolved responses to each orientation. **B**. Standard and deviant trajectories projected into the fixed, control-defined space were rotated relative to their orientation-matched controls. **C**. Diagram depicting shared adaptation (orange) resulting from repeated standard exposure (blue). **D**. Decoded stimulus angle across trials in a representative session, showing that the angular displacement emerged rapidly and remained stable within blocks. **E**. Standard and deviant responses were significantly rotated relative to held-out control responses (paired *t*-tests: C135 versus S135, *p* = 0.00377; C45 versus D45, *p* = 3.78 *×* 10^*−*6^; C45 versus S45, *p* = 0.00427; C135 versus D135, *p* = 7.36 *×* 10^*−*5^), but their rotation magnitudes did not differ significantly (block I, *p* = 0.476; block II, *p* = 0.308). Each point represents one session.

Relative to their matched control representations, both standard and deviant responses were angularly shifted on the stimulus-encoding manifold (Fig. 2B,D). These rotations occurred in the same direction and were statistically indistinguishable in magnitude (Fig. 2E). We term this “shared adaptation” because this transformation emerged rapidly following repeated exposure to the standard (Fig. 2D) and extended to both the repeated standard and orthogonal deviant representations (Fig. 2C–D). Shared adaptation changed the direction of population vectors rather than simply increasing or decreasing their magnitude. Thus, repeated exposure to a single orientation reorganized stimulus-code geometry through a shared transformation that extended beyond the repeated stimulus itself.

### Standard-specific response attenuation occurs on the stimulus-encoding manifold

Rotation changes the direction of a population vector while preserving its magnitude, whereas attenuation of responses to repeated stimuli is a well-established feature of sensory processing. We therefore asked whether presentation as a standard produced changes in response magnitude in addition to the shared rotation and whether these changes occurred on or off the stimulus-encoding manifold.

We first quantified response magnitude in the full population space and on the stimulus-encoding manifold (Fig. 3A–C). In the full space, neither standard nor deviant responses differed significantly in magnitude from their matched controls (Fig. 3B,C), although deviant responses showed a trend toward increased magnitude. On the stimulus-encoding manifold, however, standard responses were significantly attenuated relative to their matched controls, whereas deviant response magnitude did not differ significantly from control. This standard-specific reduction in on-manifold response magnitude is consistent with sensory adaptation to the repeated stimulus and represents the population component most closely corresponding to conventional SSA. Unlike shared adaptation, which alters the direction of both standard and deviant representations, this attenuation selectively affects the magnitude of the standard representation.

**Fig 3.**
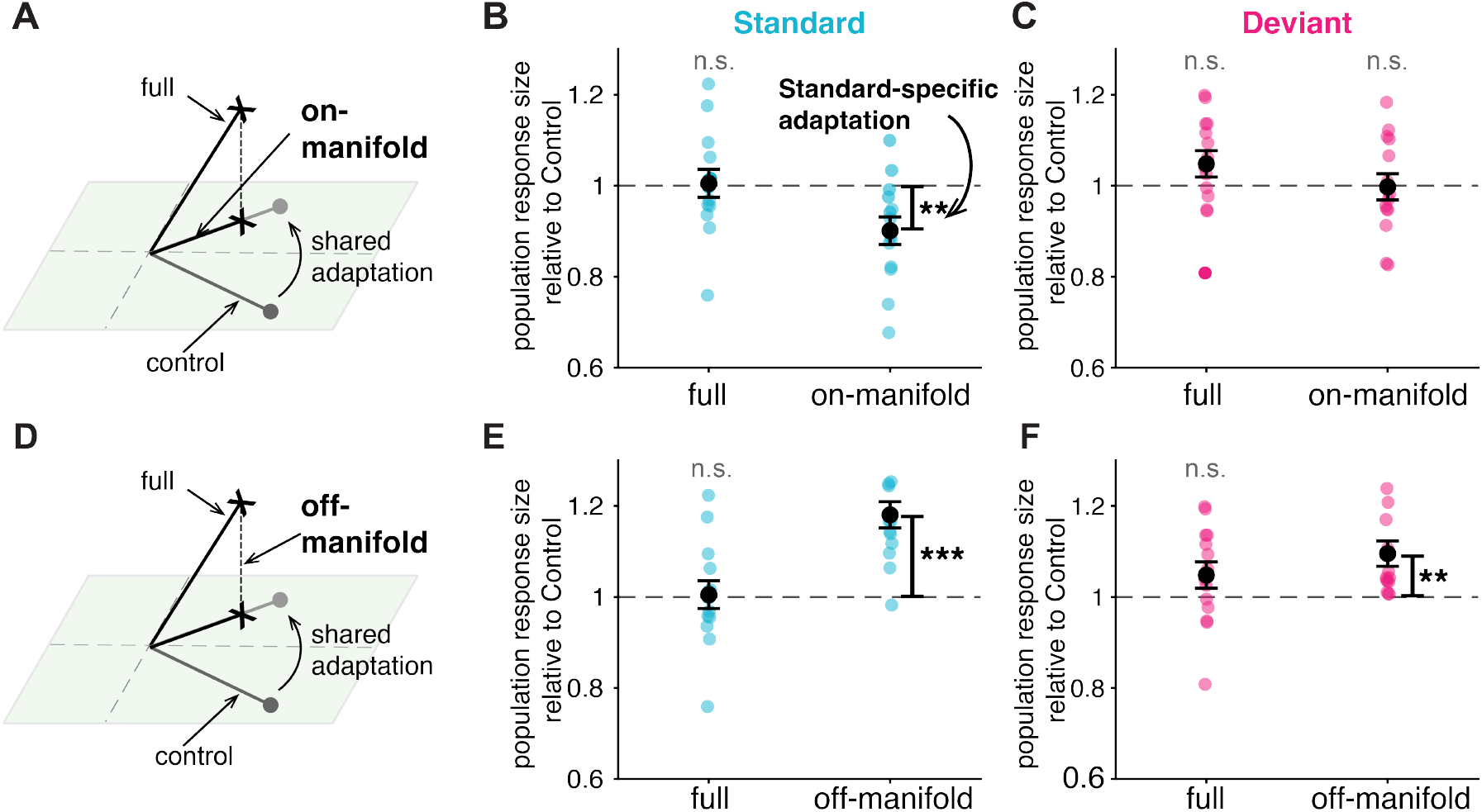
Standard-specific attenuation occurs on the stimulus-encoding manifold, whereas standards and deviants exhibit increased off-manifold magnitude. **A**. On-manifold response magnitude is the length of the projected population vector and is invariant to rotation. **B**. Standard responses did not differ significantly from controls in the full space but were reduced on the stimulus-encoding manifold relative to held-out controls (paired *t*-tests: full, *p* = 0.874; on-manifold, *p* = 0.00598). **C**. Deviant response magnitude did not differ significantly from control in either space (paired *t*-tests: full, *p* = 0.123; on-manifold, *p* = 0.923). **D**. Diagram illustrating the definition of off-manifold response magnitude. **E**,**F**. Off-manifold response magnitude was greater than that of held-out controls for both standards (**E**; *p* = 2.99 *×* 10^*−*5^) and deviants (**F**; *p* = 0.00446).

We next quantified responses in the off-manifold space, which contains patterns of activity that cannot be expressed as a linear combination of the orientation-related population patterns observed in the control condition. Both standard and deviant responses had greater off-manifold magnitude than their matched controls (Fig. 3D–F). For standards, this off-manifold increase accompanied the on-manifold attenuation, potentially accounting for the absence of a significant change in full-space response magnitude despite the reduction on the stimulus-encoding manifold. For deviants, the on-manifold component did not differ significantly from control, while the significant off-manifold increase was consistent with the trend toward increased magnitude in the full space. Standard-specific attenuation and the additional standard- and deviant-related responses therefore occupied different population dimensions, revealing geometrically separable components of visual MM. This separation raises the possibility that transformations of stimulus representations and off-manifold MM-related activity are not only geometrically distinct but also differentially dependent on cortical feedback. We therefore tested whether ACC input preferentially modulates off-manifold activity.

### ACC input selectively modulates off-manifold expectation signals

To test whether ACC feedback contributes to off-manifold MM modulation, we optogenetically inhibited ACC axon terminals in V1 during the visual oddball paradigm (Fig. 4A). Pupil size and facial motion did not differ significantly between ACC-inhibition and no-inhibition trials (Fig. S2C,D), suggesting that ACC-terminal inhibition did not produce detectable changes in these measured behavioral-state variables. We then independently defined an ACC-modulation subspace from population activity changes produced by ACC inhibition and examined its relationship with the stimulus-encoding manifold.

**Fig 4.**
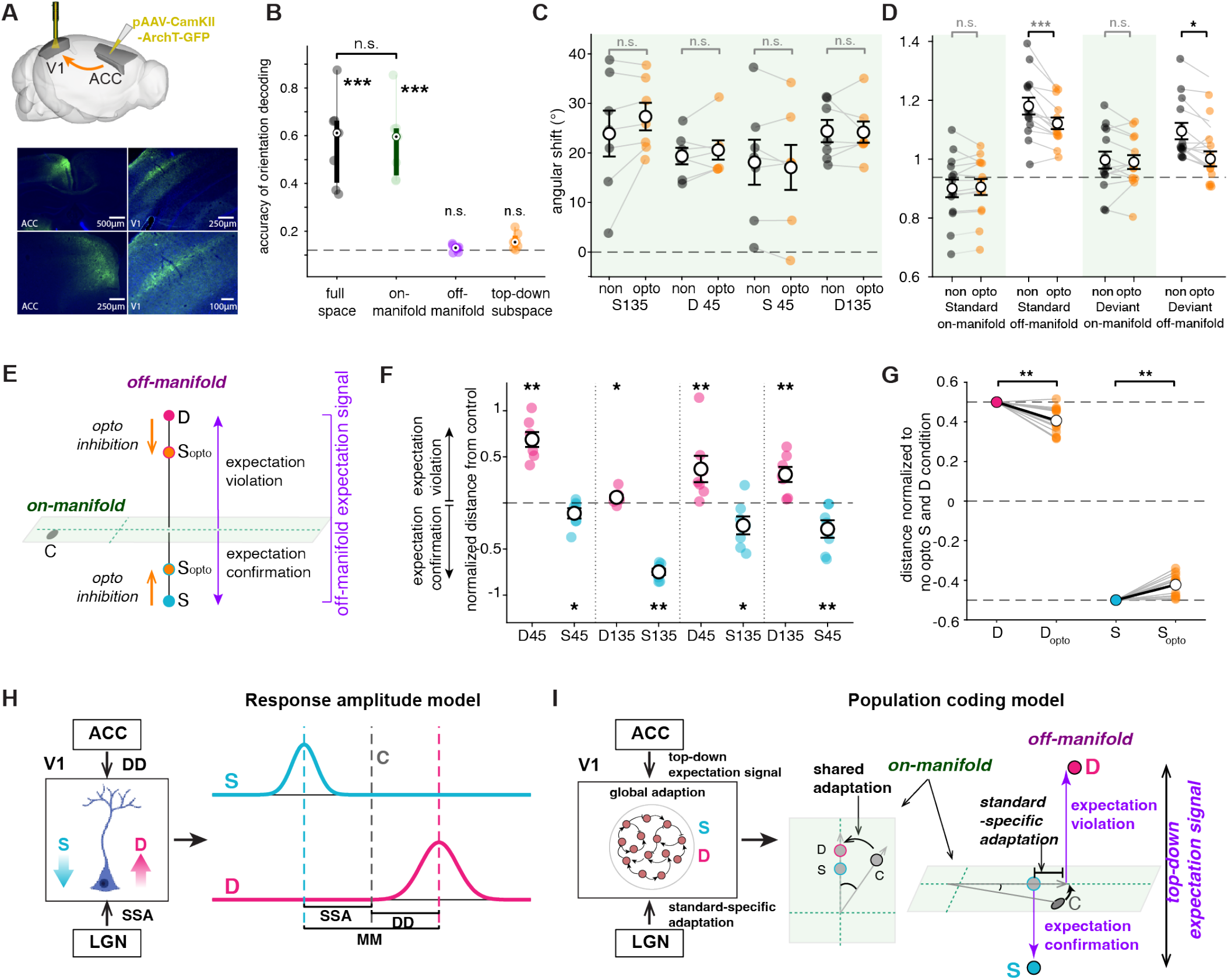
ACC input preferentially modulates expectation-related activity while preserving stimulus representations. **A**. ArchT-mediated inhibition of ACC terminals in V1 during silicon-probe recordings; images show ACC expression and projections in V1. **B**. Orientation decoding was significant in the full and on-manifold spaces but did not significantly exceed chance in the off-manifold or ACC-modulation spaces (*n* = 7 sessions; full, *p* = 5.98 *×* 10^*−*4^; on-manifold, *p* = 2.11 *×* 10^*−*4^; off-manifold, *p* = 0.305; ACC-modulation, *p* = 0.0506). Full- and on-manifold decoding were comparable (*p* = 0.934). **C**. ACC inhibition did not significantly affect standard or deviant angular rotations (paired *t*-tests: S135, *p* = 0.161; D45, *p* = 0.248; S45, *p* = 0.250; D135, *p* = 0.882). **D**. ACC inhibition reduced standard and deviant off-manifold response magnitudes (standard, *p* = 9.56 *×* 10^*−*4^; deviant, *p* = 0.0014), with a significant population-space *×* optogenetic-condition interaction (*F*_1,6_ = 7.78, *p* = 0.0316). Points represent 14 orientation–session observations from seven sessions; orientations were averaged within each session for ANOVA. **E**. Schematic of the off-manifold expectation-related axis, with standards and deviants occupying opposite poles relative to their matched controls. **F**. Standards projected toward the negative pole (S45/D45– S45, *p* = 0.0393; S135/D135–S135, *p* = 0.0076; S135/D45–S135, *p* = 0.0338; S45/D135–S45, *p* = 0.0086), whereas deviants projected toward the positive pole (D45/D45–S45, *p* = 0.0079; D135/D135–S135, *p* = 0.0244; D45/D45–S135, *p* = 0.0083; D135/D135–S45, *p* = 0.0070; paired *t*-tests, *n* = 7 sessions). These poles represent expectation confirmation and violation, respectively. **G**. ACC inhibition shifted both poles toward their matched controls (permutation tests: violation, *p* = 0.0080; confirmation, *p* = 0.0079). **H**. Conventional scalar decomposition of mismatch responses into stimulus-specific adaptation (SSA) and deviance detection (DD). **I**. Population model comprising shared on-manifold rotation, standard-specific on-manifold attenuation, and bidirectional ACC-dependent expectation-related signals off-manifold.

As expected, stimulus-orientation decoding was robust in the full population space and on the stimulus-encoding manifold (Fig. 4B) but was statistically indistinguishable from chance off-manifold. Orientation decoding was also not significantly above chance within the independently defined ACC-modulation subspace, indicating that the population dimensions modulated by ACC input carried little information about stimulus orientation.

We next asked whether ACC inhibition altered on-manifold adaptation. Shared adaptation was not significantly altered during ACC inhibition (Fig. 4C), nor was standard-specific attenuation on the stimulus-encoding manifold (Fig. 4D), suggesting that neither required ACC input.

ACC inhibition instead reduced the magnitudes of both standard and deviant off-manifold components, whereas their on-manifold components were not significantly affected (Fig. 4D). A two-factor repeated-measures ANOVA, with population space and optogenetic condition as factors, revealed a significant population-space *×* optogenetic-condition interaction, indicating that ACC inhibition affected off-manifold activity more strongly than on-manifold activity (repeated-measures ANOVA: *F*_1,6_ = 7.784, *p* = 0.0316). Thus, ACC input preferentially modulated activity outside the dimensions encoding stimulus orientation.

Response magnitude does not specify the direction in which standard and deviant responses differ from control. We therefore examined their signed geometry along an off-manifold expectation-related axis (Fig. 4E). Standards and deviants occupied opposing positions relative to their matched controls (Fig. 4F). This bidirectional geometry is consistent with the standard and deviant poles reflecting expectation confirmation and violation, respectively; the deviant-related displacement is related to the conventional DD contrast, whereas the opposing standard-related displacement reveals an additional component of mismatch processing.

Optogenetic trials were then projected onto the same fixed expectation-related axis. ACC-terminal inhibition shifted both standard and deviant responses toward their matched controls (Fig. 4G). Thus, Fig. 4D shows an unsigned reduction in off-manifold magnitude, whereas Fig. 4G reveals a corresponding contraction of both expectation-related poles toward their respective controls. ACC input therefore modulated a bidirectional expectation-related signal in off-manifold dimensions, with no significant changes detected in stimulus representations or on-manifold adaptation.

### Stimulus and expectation-related transformations occupy separable population dimensions

Together, these results reveal a population-level organization of visual mismatch in which transformations of stimulus representations and expectation-related activity occupy separable dimensions (Fig. 4H,I). Shared adaptation and standard-specific attenuation produced distinct transformations on the stimulus-encoding manifold and persisted during ACC inhibition. In contrast, standards and deviants occupied opposing positions along an off-manifold axis, consistent with expectation confirmation and violation, respectively. ACC inhibition contracted both poles toward control, demonstrating preferential modulation of this bidirectional signal by cortical feed-back. These findings identify three geometrically separable components of visual mismatch: shared adaptation, standard-specific attenuation, and bidirectional expectation-related activity. Thus, although the conventional *C − S* and *D− C* contrasts remain useful descriptive measures, neither uniquely isolates a single population-level transformation.

## Discussion

Our results reveal that visual mismatch engages separable transformations of stimulus representations and expectation-related population activity in V1. Shared adaptation and standard-specific attenuation produced distinct transformations on the stimulus-encoding manifold, whereas a bidirectional signal distinguishing standards from deviants occupied orthogonal off-manifold dimensions and was preferentially modulated by ACC input. Thus, cortical feedback preferentially modulated off-manifold mismatch-related activity, with no significant changes detected in on-manifold stimulus representations or adaptation. These population-level effects were largely obscured in average firing rates because individual units were modulated in opposing directions. Together, these findings establish a population-level framework that distinguishes shared adaptation, standard-specific attenuation, and bidirectional expectation-related activity, revealing an organization of visual mismatch that is not captured by conventional response-amplitude measures.

Shared adaptation was the more unexpected of these two transformations. Within each oddball block, standard and deviant representations rotated in the same direction and by statistically indistinguishable magnitudes relative to their matched controls. Repeated presentation of the standard therefore transformed stimulus-code geometry beyond standard-specific attenuation, with effects extending to the representation of an orthogonal stimulus presented infrequently. This shared transformation was revealed by analyzing neuronal activity in a common population state space, rather than relying on average firing rates or restricting analyses to neurons exhibiting conventionally expected response modulation. Shared adaptation extends previous evidence that orientation adaptation changes neuronal tuning and population representations in V1, including response magnitude, preferred orientation, and information available to population decoders (14, 15). Recent work has further shown that adaptation reshapes representation geometry in mouse V1 in a manner that may support efficient encoding of environmental statistics (16). Our findings extend these observations to visual mismatch processing by identifying a shared geometric transformation of standard and deviant representations. This broad generalization differs from the predominantly orientation-specific adaptation reported in primate V1, where effects are generally strongest near the adapting orientation (17, 18). Potential explanations include the locally heterogeneous organization of orientation preference in rodent V1 compared with the columnar organization of primate V1 (19, 20), or differences in the orientation selectivity of thalamic inputs (21, 22), although the present experiments do not distinguish between these possibilities.

Recurrent lateral interactions in V1 could contribute to shared adaptation, consistent with ring-attractor-like models of orientation coding (23). The confinement of shared adaptation to the stimulus-encoding manifold and its persistence during ACC-terminal inhibition are consistent with a contribution from local cortical circuits, although they do not exclude feedback from other cortical areas. Likewise, orientation-selective thalamic inputs could contribute to this transformation, leaving open the possibility of feedforward mechanisms. Notably, adaptation to either grating rotated both representations in the same direction and by statistically indistinguishable magnitudes, suggesting a common transformation that generalizes across stimulus orientations. Determining how recurrent, feedforward, and feedback circuits contribute to this shared transformation will require targeted circuit perturbations.

Population geometry separated the encoding of stimulus orientation from population activity associated with stimulus regularity and its violation. The standard–deviant axis was predominantly off-manifold, with standard and deviant responses occupying opposing positions relative to their matched controls, demonstrating bidirectional modulation. The standard-side displacement was distinct from on-manifold attenuation, consistent with an expectation-confirmation signal that is geometrically distinct from standard-specific sensory attenuation. The opposing deviant-related displacement is consistent with expectation violation and corresponds to the conventional DD contrast. Consistent with this interpretation, the independently defined ACC-modulation subspace was concentrated off-manifold, and ACC inhibition shifted both displacements toward their matched controls without significantly altering on-manifold stimulus representations or adaptation. This extends previous evidence that ACC input contributes to visual DD in V1 (24, 25) by revealing a mechanism by which ACC feedback can bidirectionally distinguish expectation confirmation and violation while fully preserving encoded stimulus orientation.

These findings also clarify the conventional *C − S* and *D − C* contrasts. At the population level, both can reflect contributions from shared adaptation, which alters both standard and deviant representations on the stimulus-encoding manifold. The *D − C* contrast also includes an off-manifold displacement consistent with expectation violation, whereas the *C − S* contrast includes an opposing off-manifold displacement consistent with expectation confirmation as well as on-manifold standard-specific attenuation. These contrasts therefore remain useful operational measurements but do not uniquely isolate the distinct population-level transformations underlying visual mismatch.

This study has several limitations. First, the many-standards control matches stimulus probability but not the full transition statistics or temporal structure of the oddball sequence. Consequently, the *D − C* contrast cannot be attributed exclusively to expectation violation. Second, our restricted stimulus set defines only the dimensions of V1 population activity associated with grating orientation. Naturalistic visual stimuli engage a richer and higher-dimensional set of sensory representations, and some activity classified here as off-manifold could therefore overlap with sensory dimensions not sampled by our stimulus set. Nevertheless, even responses to complex natural scenes occupy only a subset of the much larger V1 population space (11), leaving substantial dimensionality available for signals unrelated to stimulus identity. Determining whether these off-manifold signals remain separable from sensory representations across richer stimulus spaces will require experiments using more complex and naturalistic stimuli. Third, because animals viewed the stimuli passively, expectation confirmation and violation remain interpretations of population activity associated with sequence regularity rather than directly demonstrated behavioral expectation. Finally, most MMN studies have used auditory stimuli, and the components identified here may not generalize directly across modalities.

Together, our findings establish a population state-space framework in which stimulus representations and expectation-related activity undergo separable transformations with differential sensitivity to cortical feedback. Within this organization, visual mismatch comprises three geometrically distinct components: standard-specific attenuation and shared adaptation on the stimulus-encoding manifold, and bidirectional expectation-related modulation off-manifold. This framework identifies shared adaptation as a component not uniquely isolated by conventional SSA and DD contrasts and reframes DD as one pole of a broader signal consistent with distinguishing expectation violation from confirmation. More broadly, it provides a foundation for future work investigating how feedforward, recurrent, and feedback circuits generate these distinct population transformations. Given that reduced auditory and visual MMN responses have been investigated as a candidate biomarker of neuropsychiatric disorders (26–29), this framework may also help determine whether MMN deficits reflect widespread disruption of mismatch processing or selective impairment of its constituent population components.

## Acknowledgments

The authors wish to thank Jaslin Kalra for contributing to early stages of the project, as well as Elyse Sussman, Adam Kohn, and the members of the Batista-Brito and Sjulson labs for their helpful discussions.

## Author Contributions

R.B. and L.S. conceived the project. S.K., A.V., R.B., and L.S. designed the experiments. A.V. collected all electrophysiological data. S.K. performed all data analyses. L.S. and R.B. supervised all aspects of the project. The manuscript was written with contributions from all authors.

## Methods

### Mice

All procedures were approved by the Institutional Animal Care and Use Committee at Albert Einstein College of Medicine (protocol 00001393). Adult C57BL/6 mice were maintained on a 12-h light/dark cycle under standard housing conditions. Four mice, aged 4–7 months, were used.

### Viral injections

Mice were anesthetized with isoflurane (5% for induction, 1.5% for maintenance), placed in a stereotaxic frame, and maintained on a closed-loop heating pad. Meloxicam (2.5 mg kg^*−*1^) and topical lidocaine were administered for analgesia, and ophthalmic ointment was used to protect the eyes. A total of 500 nL of AAV-CaMKII-ArchT-GFP (Addgene catalog number 99039) was injected into each hemisphere of the ACC (AP +0.3 mm, ML ±0.25 mm, DV *−* 0.65 mm and *−* 0.75 mm relative to bregma) at 2 nL s^*−*1^ using an RWD R-480 pump and beveled glass micropipettes with 40 µm tips. A tungsten cerebellar ground wire and custom headpost were secured with OptiBond or Super-Bond and dental cement while leaving the skull over V1 accessible. Postoperative analgesia was given for 2 days, and recordings began at least 4 weeks after injection.

### Habituation and head fixation

Mice were habituated to handling for several days before surgery. After headpost implantation, mice recovered for 3–5 days before habituation to the head-fixed treadmill. Head-fixation periods were gradually increased over 1 week until animals tolerated multiple-hour sessions. The treadmill was positioned horizontally at an appropriate distance below the head and cleaned with 70% ethanol between animals.

### Electrophysiological recordings in mice

Dexamethasone (4 mg kg^*−*1^) was administered 1 h before craniotomy. Under anesthesia, a ~ 3 mm craniotomy was made over V1 and covered with Kwik-Cast silicone elastomer (World Precision Instruments). Mice were allowed to recover for at least 1 h before recording. A 64-channel linear silicon probe (H3, Cambridge NeuroTech) coupled to a tapered optical fiber was inserted into V1 at 1 µms^*−*1^ over ~ 20 min. Silicone oil was used to prevent drying of the craniotomy. After each session, the craniotomy was resealed with Kwik-Cast; mice were recorded once daily for 3–4 days. Blue light was delivered through the fiber using an LED (MF470F4, Thorlabs) controlled by custom MATLAB software, an NI PCIe-6323 interface, and an LEDD1B driver (Thorlabs). No photoelectric artifacts were detected. Signals were acquired at 20 kHz using an Intan RHD2000 interface.

### Visual stimulus protocol

Recording sessions consisted, in order, of a 15-min spontaneous-activity block with a gray screen, followed by three repetitions of a sequence comprising two oddball blocks followed by one control block. Thus, each session contained six oddball blocks arranged as three flip-flop pairs and three control blocks. A 2-min interval separated consecutive blocks. Drifting gratings were presented for 500 ms. The nominal interstimulus interval was 700 ms. Gratings had 100% contrast, a spatial frequency of 0.2 cycles/cm, and a temporal frequency of 2 Hz. Stimuli were generated in MATLAB using Psychtoolbox.

In each oddball block, the 45° and 135° gratings served as the standard and deviant orientations. Each block contained 768 stimulus presentations: 672 standards and 96 deviants. Thus, standards and deviants accounted for 87.5% and 12.5% of presentations, respectively. The 45° and 135° gratings exchanged standard and deviant roles between the two blocks of each flip-flop pair (blocks I and II). Across a complete session, three blocks used each orientation as the standard and three used it as the deviant. In the control block, eight orientations—0°, 22.5°, 45°, 67.5°, 90°, 112.5°, 135°, and 157.5°—were each presented 80 times, producing 640 trials per block. The 640 presentations were randomly permuted. Optogenetic inhibition of ACC terminals in V1 was assigned separately to standard and deviant presentations. Within each oddball block, half of the deviants (48 per block) were randomly selected for optogenetic inhibition. In addition, 48 standard presentations were randomly selected for optogenetic inhibition in each block. In the control blocks, half of the presentations of the two oddball-associated orientations, 45° and 135°, were randomly selected for illumination; the other six orientations were not illuminated. The laser was turned on 100 ms before stimulus onset and turned off at stimulus offset.

### Histology

Histology confirmed viral expression in ACC and labeled ACC axons in V1. Mice were perfused with 4% paraformaldehyde in phosphate-buffered saline (PBS), and 20 µm coronal sections were prepared using a CryoStar NX70 cryostat. Sections were blocked for 1 h at room temperature in 1.5% normal donkey serum and 0.1% Triton X-100 in PBS, incubated with rat anti-GFP primary antibody for 48 h at 4 °C, and then incubated with donkey anti-rat secondary antibody for 1 h at room temperature. Slides were coverslipped with DAPI-containing mounting medium (ProLong Gold, Invitrogen, P36935) and imaged on a Zeiss AXIO Imager.A1 microscope using 10*×* or 20*×* objectives.

## Data analysis

### LFP and unit analysis

LFP data were obtained from the broadband recording sampled at 20 kHz by independently applying a 200 Hz Butterworth low-pass filter to each channel and downsampling to 2 kHz. For subsequent analysis, each channel was z-scored. Trials were separated according to stimulus orientation (45° or 135°) and sequence condition (standard, deviant, or control). Only trials without optogenetic illumination were included in this analysis. Channels identified as bad were excluded. For each session, a representative channel was selected as the channel showing the largest negative response, averaged across standard, deviant, and control trials at both orientations, during the 0–100-ms post-stimulus interval. LFP responses were averaged across trials, and response magnitude was quantified as the area under the trial-averaged waveform during the 0–100-ms poststimulus interval.

Single units were isolated using Kilosort2.5 with default parameters, followed by manual curation in Phy to remove noise clusters, considering refractory-period violations, waveform shapes, and firing-rate stability. Both single and multiunits were included in the analysis. Spike counts were measured during the 500 ms stimulus presentation. Trial numbers for each orientation were balanced across standard, deviant, and control conditions by randomly subsampling each condition to the smallest trial count. Units were classified as positively or negatively modulated according to the sign of the relevant contrast, *C − S, D − C*, or *D − S*.

### Population decoding

Standard and deviant trials were classified separately for each orientation and session using linear support vector machines implemented with MAT-LAB’s `fitcecoc`. Inputs were unit spike counts during the 500 ms stimulus presentation. Aggregate activity was represented by the mean firing rate across units. Accuracy was evaluated using fivefold cross-validation repeated 10 times. Within each fold, features were z-scored using training-set statistics. Decoding was performed using aggregate activity, the full population, and MM-positive and MM-negative subpopulations. Subpopulations were identified within each fold using only training trials. Units were classified using two-sided Welch’s *t*-tests comparing standard and deviant responses (*p <* 0.025). For each fold, a shuffled baseline was generated from 100 permutations of the training labels while retaining the true test labels. The accuracy index was the observed accuracy minus the mean shuffled accuracy.

### Construction of the stimulus-encoding manifold

The stimulus-encoding manifold was constructed separately for each recording session from spike counts measured during the 500 ms stimulus presentation. Spike counts were standardized independently for each unit across the control trials and then averaged across trials of the same orientation, producing an 8-orientation *× N*-unit matrix. After centering across orientations, principal component analysis was applied and the first seven dimensions were retained. Standard, deviant, and control responses were projected into this stimulus-encoding manifold, and off-manifold activity was defined as the orthogonal residual.

### Shared adaptation analysis

Shared adaptation was quantified within a seven-dimensional stimulus-encoding manifold as the signed difference between the presented orientation and the orientation decoded from the population response. A circular ridge-regression decoder was trained on control-trial responses to predict double-angle coordinates, [cos(2*θ*), sin(2*θ*)], accounting for the 180° periodicity of orientation. The signed angular shift was calculated as

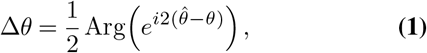

where *θ* is the presented orientation and 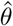 is the decoded orientation. Rotations were therefore restricted to *−* 90° to 90°, with positive and negative values indicating opposite directions of rotation within the stimulus-encoding manifold. Decoders were fitted using a fivefold partition of control trials and applied to standard and deviant population responses. The analysis was repeated 100 times with randomized control-trial sampling and fold assignment. Angular rotations were averaged across folds and repetitions. Standard and deviant rotations were compared across sessions using two-sided paired *t*-tests.

### Population-response magnitude on and off the stimulus-encoding manifold

For each trial, the population response vector was constructed from spike counts during the 500 ms stimulus presentation. For each unit, responses were z-scored using the mean and standard deviation of its responses on control trials. For each orientation, on-manifold responses were obtained by projection onto the seven-dimensional stimulus-encoding manifold, whereas off-manifold responses were defined as the residual after removing this projection. Population response size was calculated for each trial as the Euclidean norm of the full-space, on-manifold, and off-manifold response vectors. To account for general response attenuation, the off-manifold magnitudes were divided by the corresponding normalized on-manifold magnitude separately for each orientation. For each orientation, standard and deviant magnitudes were divided by the mean magnitude of the corresponding control responses, such that a value of 1 represented the mean control response. Balanced trial selection was repeated 10 times. Each orientation–session pair was treated as one observation, and group summaries show the mean *±* SEM. Normalized population response size was compared with the control value of 1 using a two-sided one-sample *t*-test.

### Off-manifold expectation-related axes and ACC modulation

Off-manifold expectation-related signals were quantified from population spike counts during the 500 ms stimulus period. For each session, an expectation-related axis was defined as the orthogonal residual of the no-opto *D − S* population vector after removing its projection onto the stimulus-encoding manifold. Axis signs were oriented so that deviant projections were greater than standard projections, with control centered at zero. Axes were constructed for orientation-matched comparisons (*D*_45_ *− S*_45_ and *D*_135_ *− S*_135_) and within-block comparisons (*D*_45_ *− S*_135_ and *D*_135_ *− S*_45_). Fivefold cross-validation was used; axes, signs, and normalization were determined from training trials and applied to held-out test trials. Projections were centered on the control response and normalized by the training-set *D − S* distance. Deviant projections above control and standard projections below control were evaluated across sessions using one-sided sign-flip permutation tests with 100,000 permutations.

The effect of ACC inhibition was measured along orientation-matched off-manifold expectation-related axes. No-opto trials defined the axis, its sign, and its normalization. ACC-inhibited standard and deviant responses were projected onto the fixed no-opto axis and expressed relative to the no-opto control response. Projection distances were normalized by the no-opto *D − S* separation. For visualization, no-opto *D* and *S* positions were fixed at *−* 0.5 and 0.5, respectively. ACC-inhibited responses were displaced according to the change in their distance from the opposite no-opto response. Thus, movement toward zero indicated that ACC inhibition shifted activity toward the opposite no-opto endpoint. For each session, results were averaged across the two orientations. Optogenetically induced changes in distance were evaluated across sessions using one-sided sign-flip permutation tests with 100,000 permutations; negative distance changes indicated movement toward the opposite no-opto response.

### Pupil and behavioral-state analysis

Videos of the mice were acquired at 30 Hz using Blackfly cameras (FLIR) controlled by FlyCapture2 software. Camera-generated TTL pulses were recorded with the electrophysiological data to synchronize video frames with neural recordings. Pupil size and facial motion energy were extracted from each video frame using FaceMap software.

### Statistical analysis

Statistical analyses were performed in MATLAB, with significance defined as *p <* 0.05. Unless otherwise specified, tests were two-sided, and data are reported as mean *±* SEM. Paired *t*-tests were used for paired comparisons, and normalized response magnitudes were compared with the control value of 1 using a one-sample *t*test. Decoding accuracy indices were compared with zero using one-sample *t*-tests. Directional effects of off-manifold expectation-related signals and ACC-induced changes were evaluated using one-sided sign-flip permutation tests. Selective off-manifold modulation was tested using a two-factor repeated-measures ANOVA with population space and optogenetic condition as within-session factors.

## Declaration of generative AI and AI-assisted technologies in the manuscript preparation process

During the preparation of this work, the authors used Chat-GPT to assist with language editing and improving the clarity of the manuscript. The authors reviewed and edited the output as needed and take full responsibility for the content of the published article.

