## Supplementary Information for "Distinct population dimensions separate sensory adaptation and expectation-related signals in visual cortex"

Distinct population transformations underlie visual mismatch responses in V1

### Supplementary Figures and Legends

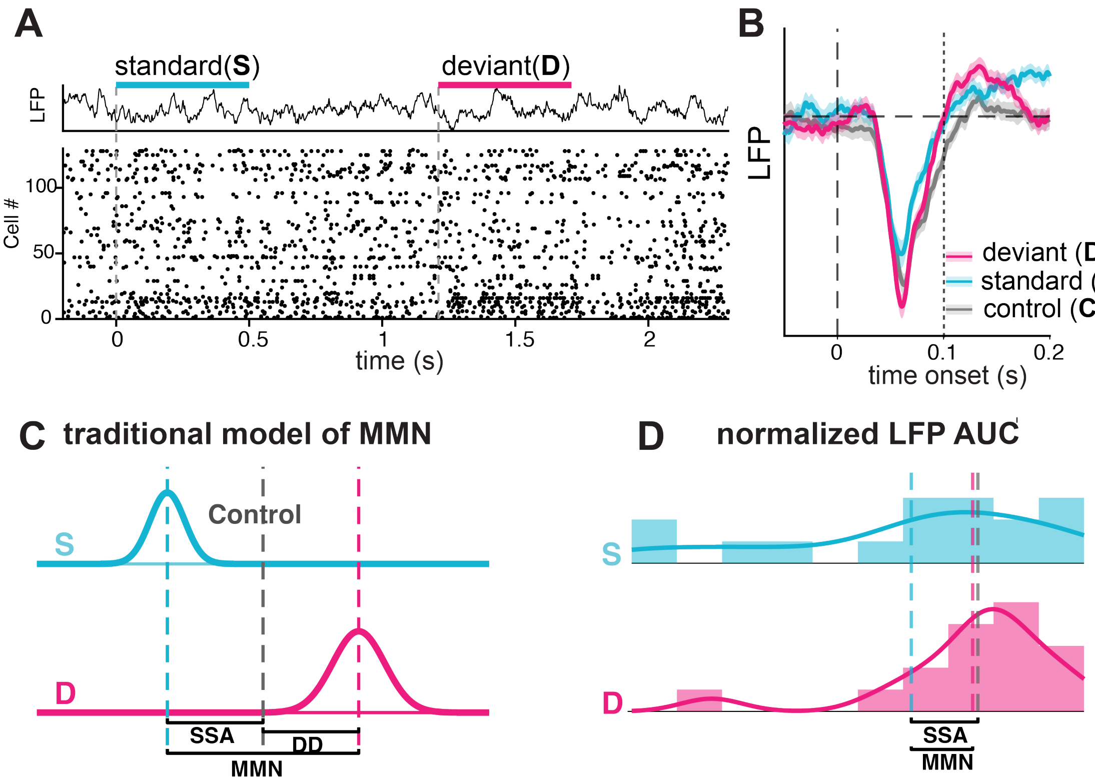


**Figure S1. LFP responses provide an aggregate reference for visual mismatch. A.** Conventional scalar model in which $C-S$ is termed stimulus-specific adaptation (SSA), $D-C$ is termed deviance detection (DD), and $D-S$ is the mismatch (MM) contrast. **B.** Normalized LFP response magnitude for standard, control, and deviant stimuli. **C.** Example standard and deviant LFP time courses. **D.** Summary of LFP responses. LFP measurements capture aggregate differences among presentation conditions but do not resolve the population geometry examined in the main figures.

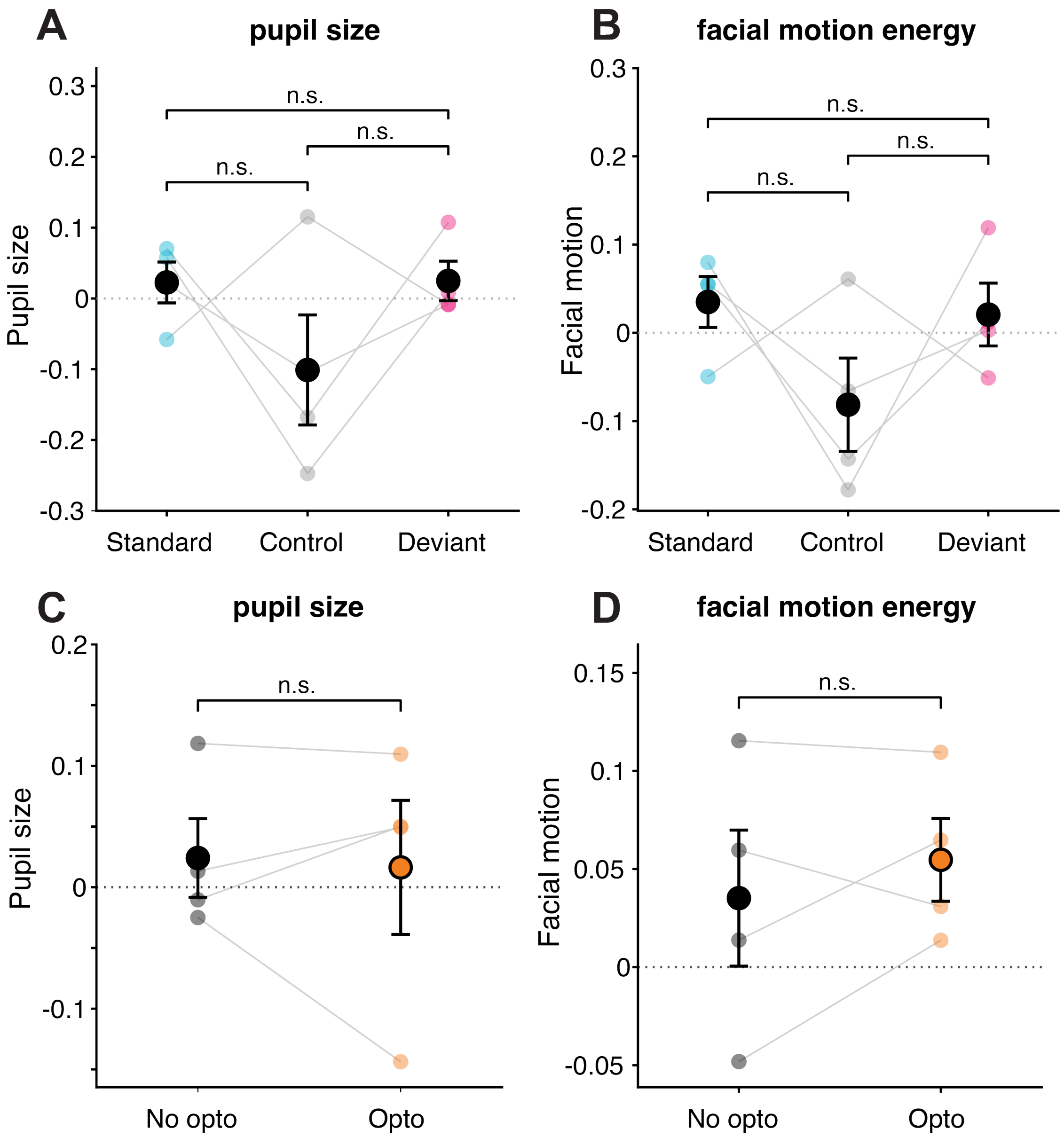


**Figure S2. Measured behavioral state across visual-stimulus and optogenetic conditions. A,B.** Trial-averaged pupil size (**A**) and facial motion energy (**B**) during standard, control, and deviant trials without optogenetic illumination. No significant differences were detected among stimulus conditions ($n = 4$ mice; two-sided paired tests: pupil size, S versus C, $p = 0.326$; C versus D, $p = 0.267$; S versus D, $p = 0.932$; facial motion energy, S versus C, $p = 0.245$; C versus D, $p = 0.318$; S versus D, $p = 0.547$). **C,D.** Pupil size (**C**) and facial motion energy (**D**) during ACC-terminal-inhibition and no-inhibition trials in the oddball paradigm. Neither measure differed significantly between optogenetic conditions ($n = 4$ mice; two-sided paired tests: pupil size, $p = 0.195$; facial motion energy, $p = 0.438$). Points indicate individual mice; summary values indicate mean ± SEM.
